# A structural approach to disentangle the visualization of bipartite biological networks

**DOI:** 10.1101/192013

**Authors:** J. Garcia-Algarra, J. M. Pastor, M. L. Mouronte, J. Galeano

**Affiliations:** Centro Universitario U-TAD, Las Rozas, Spain; Complex Systems Group, Universidad Politecnica de Madrid, Spain; ETSIAAB, Dept. Ingenieria Agroforestal, Universidad Politecnica de Madrid, Spain; Computer Science Department, Universidad Francisco de Vitoria, Madrid, Spain

**Author notes:** Corresponding author: Email addresses (J. Galeano).

**Keywords:** Biological bipartite networks, visualization, *k-core* decomposition

## Abstract

Interactions between two different guilds of entities are pervasive in biology. They may happen at molecular level, like in a diseasome, or amongst individuals linked by biotic relationships, such as mutualism or parasitism. These sets of interactions are complex bipartite networks. Visualization is a powerful tool to explore and analyse them but the most common plots, the bipartite graph and the interaction matrix, become rather confusing when working with real biological networks. We have developed two new types of visualization that exploit the structural properties of these networks to improve readability. A technique called *k-core decomposition* identifies groups of nodes that share connectivity properties. With the results of this analysis it is possible to build a plot based on information reduction (Polar Plot) and another which takes the groups as elementary blocks for spatial distribution (Ziggurat plot). We describe the applications of both plots and the software to create them.

## 1. Introduction

Network Science is a powerful tool for biological research across all scales: molecular [1, 2, 3], genetic [4, 5, 6], individual [7, 8] and communitary [9, 10]. The conceptual framework is valid for them all, and this fact has fostered both theoretical and applied developments. An important subset of biological networks are bipartite. They have two different classes of nodes. Each one that may be tied to nodes of the opposite guild but never to its peers.

Gene-protein, host-pathogen and predator-prey interactions are the basis of bipartite biological networks. A common structural property of them is the core-periphery organisation [11, 12, 13]. This fact is well-known in ecology. In mutualistic communities there is one group of very interconnected nodes, the `generalists`, that provide stability and resilience [14]. Species with a low number of links (`degree`) are tied to those specialists. This property is called nestedness and there are different indexes to measure its strength [15]. Another important structural feature is modularity, that accounts for the existence of small groups of nodes of high degree inside a network sparsely connected [16].

In many cases the issue of interest is not the generalization of the network properties but the study of a particular system itself. In these fields dealing with complex systems scientists are more interested in finding special relationships or understanding the role of a specific node than their statistical properties. A more detailed, qualitative rather than quantitative analysis about relationships in a complex network may be more useful for some researchers in Medicine, Biology, Sociology or even Economy. Visualization may play an important role in network analysis as it between data and people [17, 18, 19].

The range of possible applications is wide [20]. For instance, a field ecologist could identify central species and those most endangered within a community with a good network plot. A clinical researcher may detect anomalies in complex gene-protein associations. Visualization is an essential procedure in the exploratory stage [21], but it requires fast and interactive applications able to disentangle structure. Although a lot of effort has been put in analytic tools development, those designed for bipartite biological visualization are still scarce [22, 23, 24, 25, 26, 27].

The most common plots in literature are the bipartite graph and the interaction matrix, two ways to visualize a bipartite network of any kind. In the *bipartite graph*, nodes of both classes are plotted along parallel lines. Interactions appear as links amongst them (fig. 1). On one hand, it is quite simple, as it makes clear the separation of guilds. On the other, it is not easy to follow indirect interactions, those between two nodes of the same class linked by a common node of the opposite. They are not much relevant in affiliation networks (journals-authors, movies-actors) [29], but are extremely important in many biological networks. They create feedback loops that increase complexity and eventually emerging properties that arise from it [30, 31].

**Figure 1:**
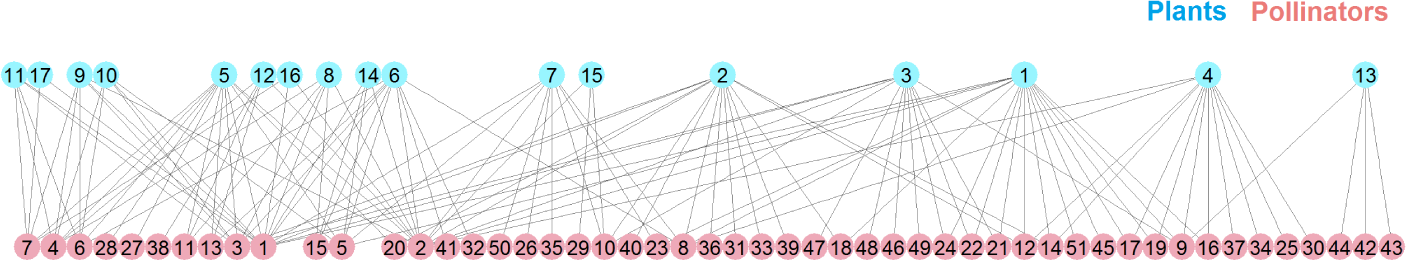
Mutualistic community in Tenerife, Canary Islands (Spain), with 68 species and 129 links [28]. In mutualism, species fall into two disjoint guilds, such as plants and pollinators or plants and seed dispersers. Ties amongst species of the same guild are forbidden.

When the number of nodes of a bipartite plot is in the range of a few tens it becomes extremely confusing. It is hard to distinguish individual links and impossible to follow indirect interactions. Accumulation of links in the space between guilds creates what is known as the *hairball effect* [32], but the main shortcoming of the bipartite plot is that it does not show the network hyerarchical organisation.

In the *interaction matrix*, nodes of one guild are arranged along rows and species of the opposite guild along columns. A filled cell marks the interaction between two species. With the interaction matrix it is possible to visually discover patterns of nestedness and modularity, so it is more expressive in the representation of structure. On the other hand, indirect interactions are even less apparent than in the bipartite plot. The matrix also becomes difficult to interpret when the number of nodes and links raise.

To overcome the drawbacks of the bipartite graph and the interaction matrix there are two possible attack strategies: information reduction or taking advantage of known network traits to order nodes and links in space. In this paper, we explain how structural properties of bipartite biological networks are the basis of two new types of visualization. Both rely on a classical technique called *k-core decomposition*. We also describe an interactive application to plot them.

## 2. Plots

The rationale behind this research is that, as biological networks are not random, this fact should provide a natural way to group nodes using their topological properties. These groups must be the basis for a spatial distribution that minimizes the hairball effect and, in addition, makes structural sense.

The *k-core decomposition* is a fast and efficient technique to cluster nodes by their connectivity properties [33, 34]. The *k-core* of a graph *G* is a maximal connected subgraph of degree *k*. Each node of the core of order *m* (called *m-shell*) has links with at least *m* other nodes that belong to that same core. The practical implication of this definition is that nodes are classified according to their connectivity. The innermost shell is the set of highest *k* index nodes. Nodes with higher degrees are the *generalists*. As *k* index decreases, nodes become more specialist. The usual way to identify the *m* - *shell* subsets is the *pruning* algorithm: one starts pruning the nodes with just one link, recursively. This subset of nodes constitutes the 1-*shell*. The remaining nodes are tied by at least two links. In the next step one extracts nodes with only two links, also recursively; this subset is the 2-*shell*. And so on. This procedure helps to recognize how the nodes of the m-shell are tied to the network. We refer to [35] for further details on the k-core analysis of bipartite networks.

As a result of the analysis we define two magnitudes. The first one is *k_radius_*. The 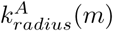 of node *m* of guild *A* is the average distance to all nodes of the innermost shell of guild *B* (set *C^B^*).

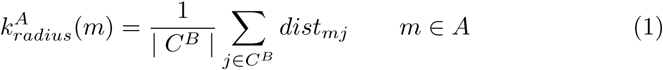

where *dist_mj_* is the shortest path from node *m* of guild *A* to node *j* of guild *B*. In an intuitive way, *k_radius_* measures how far the node is from the most connected shell, the group that is the corner stone of the network; *the closest to one the strongest the tie to the core*.

The second magnitude is *k_degree_*. It is defined as the sum of the inverses of *k_kradius_* of neighbour nodes:

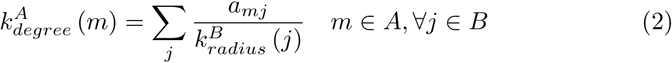

where *a_mj_* is the element of the interaction matrix that represents the link, considered as binary (1 if *a_mj_* > 0, 0 if *a_mj_* = 0). Note that this magnitude is a weighted degree where the weight is the inverse of the *k_radius_*, in such a way that links to the periphery are underestimated, so *k_degree_* is a measure of centrality.

### 2.1. The Polar plot

The *k-core* decomposition helps to visualize very large systems and networks and to understand their structure [36, 37]. In particular, the *fingerprint plot*, uses a polar coordinate system [38]. Nodes are depicted at a distance pro-portional to the shell they belong to and their areas are proportional to their `degree`.

Taking this idea as the starting point, we build the *polar plot*. Differences are noteworthy. The first one is the bipartite nature of the networks, so space is divided in two half planes, one for each guild. Node shapes are also different for each guild. This plot provides an overview of how far from the core the nodes are and, at the same time, their connectivity (by the size of the marker) and to which *m* - *shell* they belong (by the color of the marker). This visualization is interesting to detect some special features of the network; for instance, a well bonded core will present the innermost shell at distance *k_radius_* equals to one, and a nested network will show a periphery close to the core. This plot shows the periphery nodes less relevant for the network connectivity as markers far away from the core and it allows to detect highly connected nodes that do not belong to the core. Angle does not convey information, the algorithm computes it to reduce node overlapping. Links are not displayed.

Optionally, the user may choose to display the histograms of *k_degree_*, *k_radius_* and *k*-*shell*. The *k_radius_* histogram shows the distribution of node distances to network core. The *k_degree_* histogram is very similar to the degree distribution but with non-integer bins, due to the weights in its definition. The most interesting histogram is that of the *k_shell_*; a typical nested network exhibits a U-shaped *k_shell_* histogram. This shape of distribution is related to a big core and numerous peripheral nodes; a L-shaped histogram is related to a network with too many peripheral nodes and a small core.

**Figure 2.**
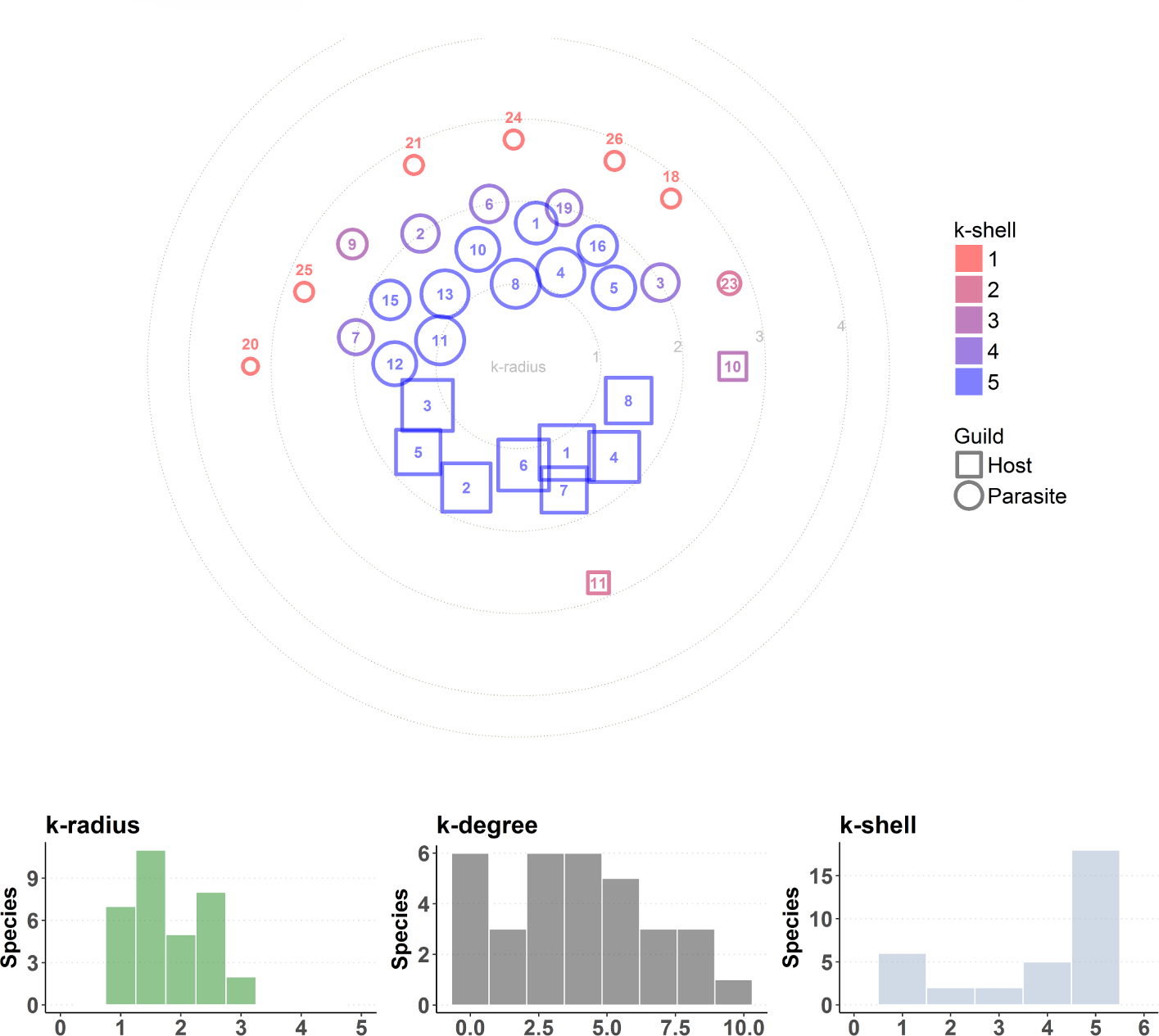
Host-parasite assembly in Tyva (Russian Federation) [39].

Figure 2 is the polar plot of a host-parasite assembly with a characteristic high concentration of nodes in the innermost shell. Most nodes lay inside the *k_radius_* 1 circle but there is a sensible number of outlying species. This network is moderately nested (*NODF* = 29).

### 2.2 The Ziggurat plot

The polar plot does not show network links, as it works on the information reduction strategy. The ultimate goal of this research is the creation of a new kind of diagram with as many details as possible. The basic idea is grouping nodes by their *k* - *shell*. If we stick nodes with the same *k* - *index* in a reduced area, links amongst the same shell nodes will not spread across the whole space. Only ties with their edges in different shells would have long paths. This simple principle is not so easy to implement. The bipartite nature of networks means that links have to go from one guild to the opposite.

The core-periphery organization implies that there are many ties from 1-*shell* nodes towards upper *k* - *index* groups. Nodes with high degree are prone to be visually suffocated by surrounding links in the bipartite graph. See plant species numbers 1, 2, 3, 4 in figure 1. This danger is a formidable obstacle.

Figure 3 shows a ziggurat plot under construction. It is the same network of fig. 1. The *k-core* decomposition puts each species inside one shell, we do not show nodes of 1-*shell* at this moment. The maximum *k* - *index* is 4 for this community.

**Figure 3:**
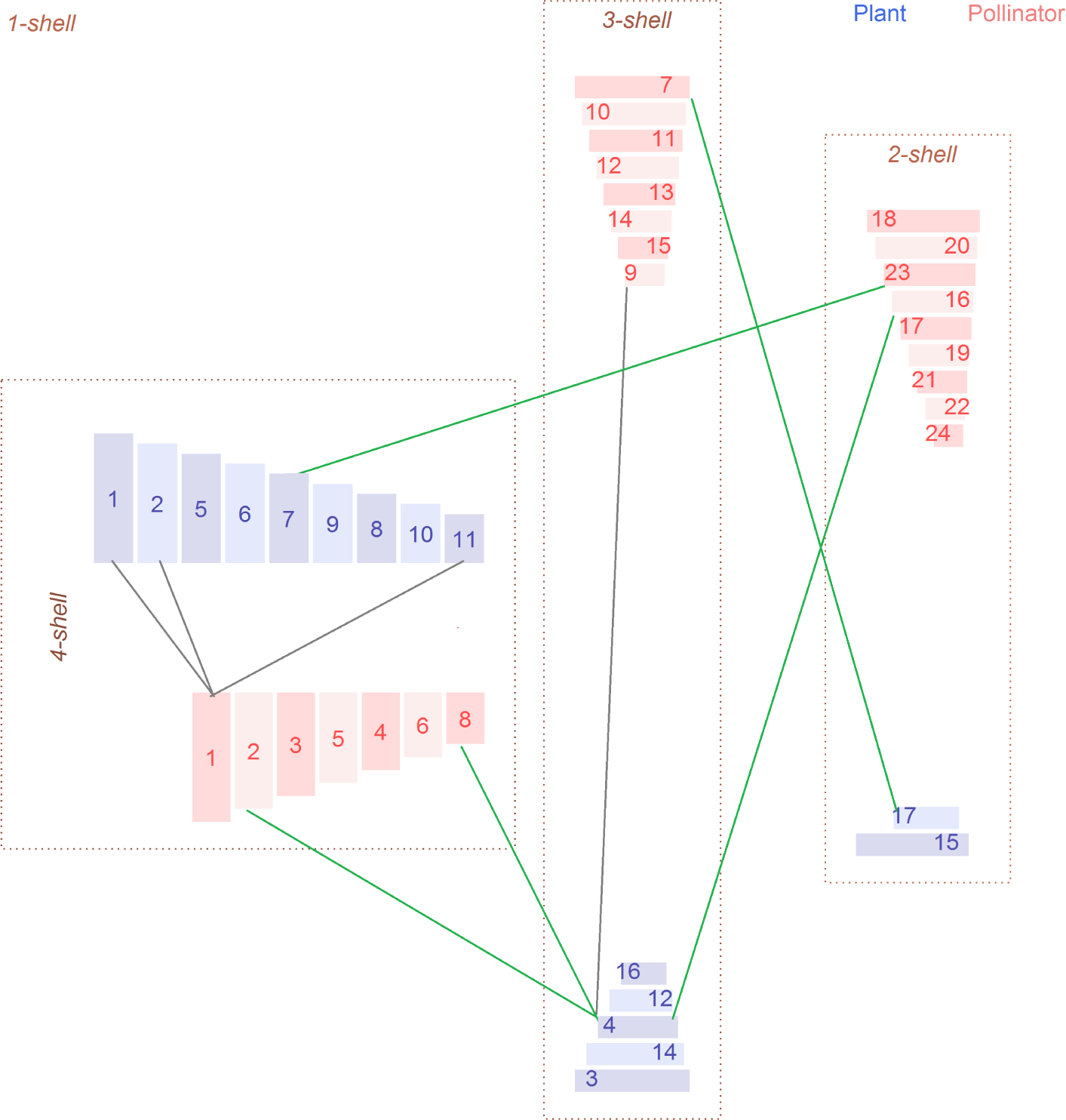
Building blocks of the ziggurat plot.

The innermost shell is found on the center of the plot, slightly leftwards. Nodes are rectangular-shaped, and are ordered by *k_degree_*. Heights decrease just for plotting convenience. The specular position of both guilds leaves space to draw the links amongst them. In fig. 3 we have plotted just three connections from *pollinator* 1 towards plants of 4-*shell*.

Lower *k* - *shells* have ziggurat shape, with nodes ordered by ascending *k_radius_*, so *pollinator* 7 is the closest to the innermost shell in 3-*shell*. Links inside the shell (gray color) connect the left sides of rectangles (*plant* 4-*pollinator* 9). Links between two different shells (green) connect the right side of the highest *k* - *index* node to the left side of the lowest one (*plant* 17-*pollinator* 7).

3-*shell* ziggurats are more distant from the horizontal axis than 2-*shell* ziggurats. Moving them up or down, it is possible to change the area of the internal almond-shaped space defined by the ziggurats and the innermost shell triangles. This area is key because links from 4-*shell* lay here, and do not cross the inner ziggurats.

The outer space is the 1-*shell* nodes home. We divide them into three groups: outsiders, tails and chains of specialists. Outsiders are nodes disconnected from the giant component. They are unusual in recorded ecological networks because by definition they do not interact with the community. This network lacks outsiders. Tails are nodes directly connected to higher*k*-*indexnodes*. They are very common, and to reduce the number of lines we apply a simple grouping rule. If *n* tails are tied to the same species of a ziggurat, we plot them in a unique box with just one link. Chains of specialists are less frequent. They are built with nodes of 1-*shell* linked amongst them, the edge that has a link with a higher*k*-*index* shell is the root node (*plant* 13).

Putting everything together, we obtain the ziggurat plot of fig. 4. This may be compared with the bipartite graph of the same network (fig. 1).

**Figure 4:**
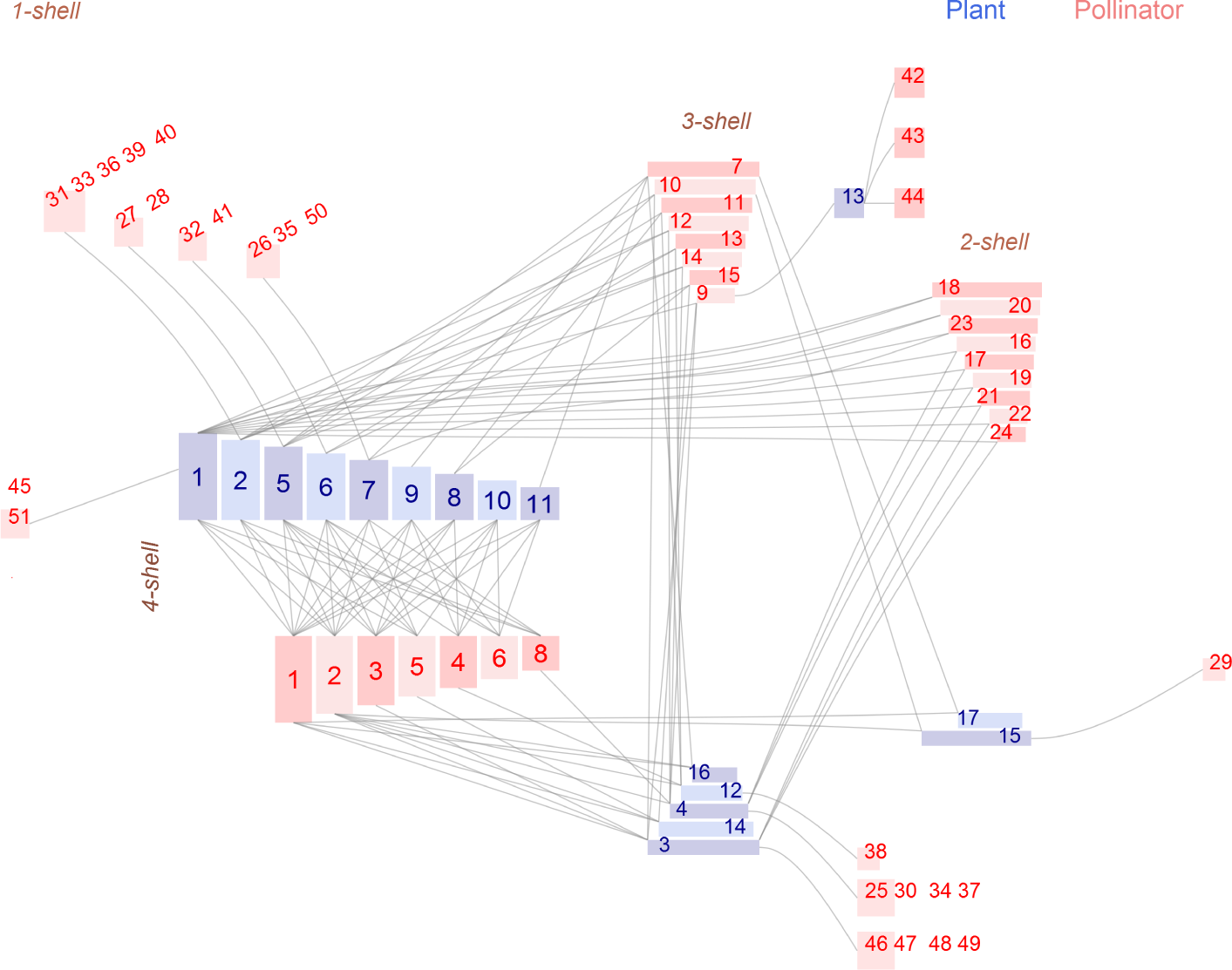
Ziggurat plot of the same plant-pollinator community in Tenerife, Canary Islands (Spain), that appears in fig. 1.

Links are drawn as straight lines or splines, that make the diagram more appealing to the eye. If links are weighted, setting the width of each link to be proportional to a function of the interaction strength is optional.

*Note that, for a given node, the main links are those towards higher k-shells. With this plot is very easy to observe how many links depart from a node to higher and to lower k-shells. One can also compare if a node is connected to higher k-shell than another one, and then if its contribution to the network is more important*.

## 3. Exploratory analysis using *k-core* plots

- The ziggurat plot unveils structural details that are hard to visualize in the bipartite graph. Figure 5 is a network of associations amongst human diseases and non-coding RNA (IncRNA), we refer to the original paper to compare with the bipartite visualization [40]. It is a small network with 39 nodes and low connectivity, just 44 links. The highest degree IncRNA node is number 8 (XIST), that in the bipartite plot looks as the most central one. The ziggurat shows at a glance that despite its high connectivity, it appears in association with diseases that belong to its chain of specialists. On the other hand, diseases like breast cancer and acute myeloid leukemia are associated with multiple *IncRNAs*.

**Figure 5:**
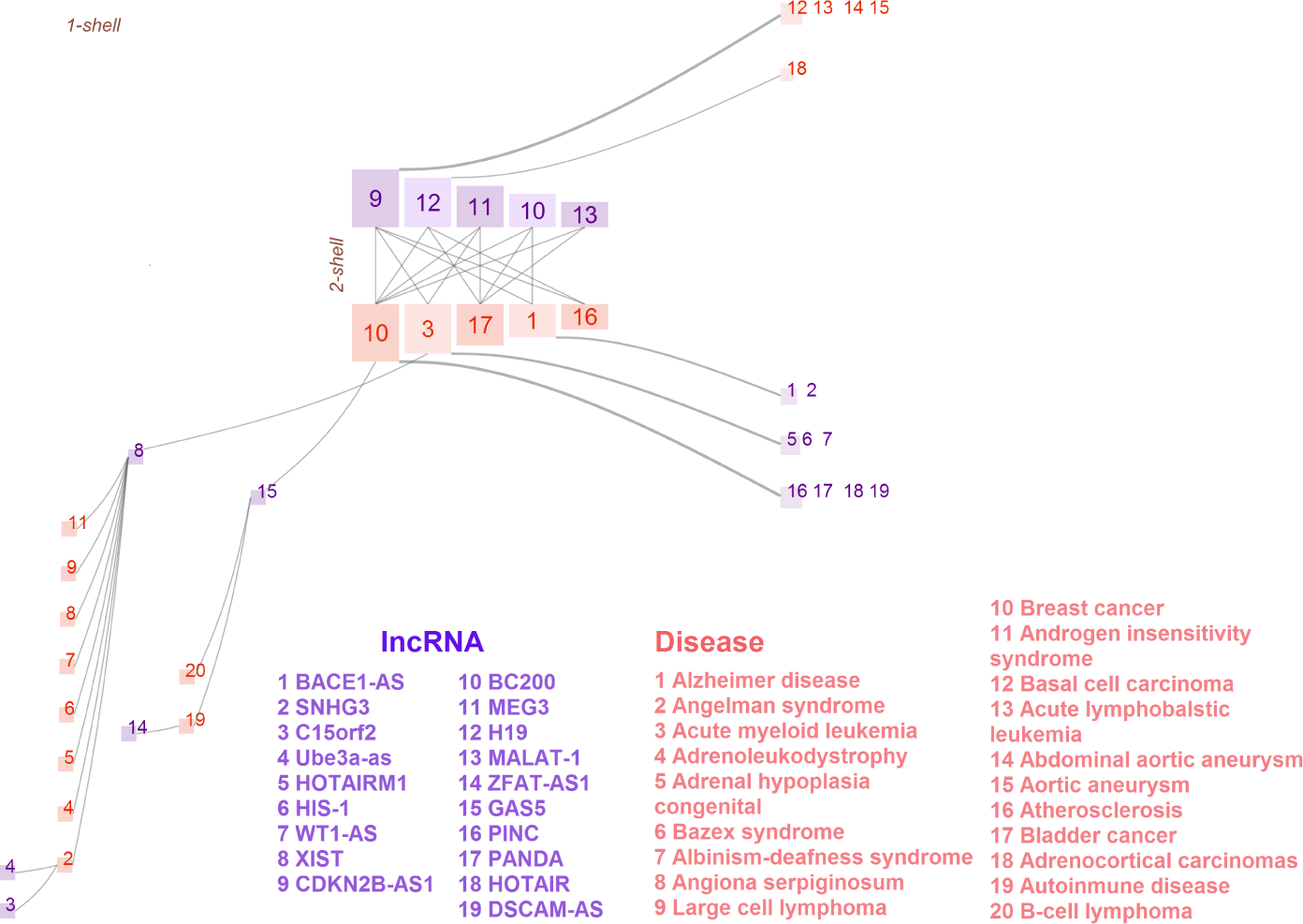
IncRNA-disease association network [40].

The network of figure 6 is slightly bigger, with 29 gene signatures used for predicting the reoperative treatment response of breast cancer and 19 pathways to different types of cancer [41]. The bipartite plot is hard to understand in the original paper, because of the number of ties. With 149 links, it becomes a *hairball* and the problem is just its own nature.

**Figure 6:**
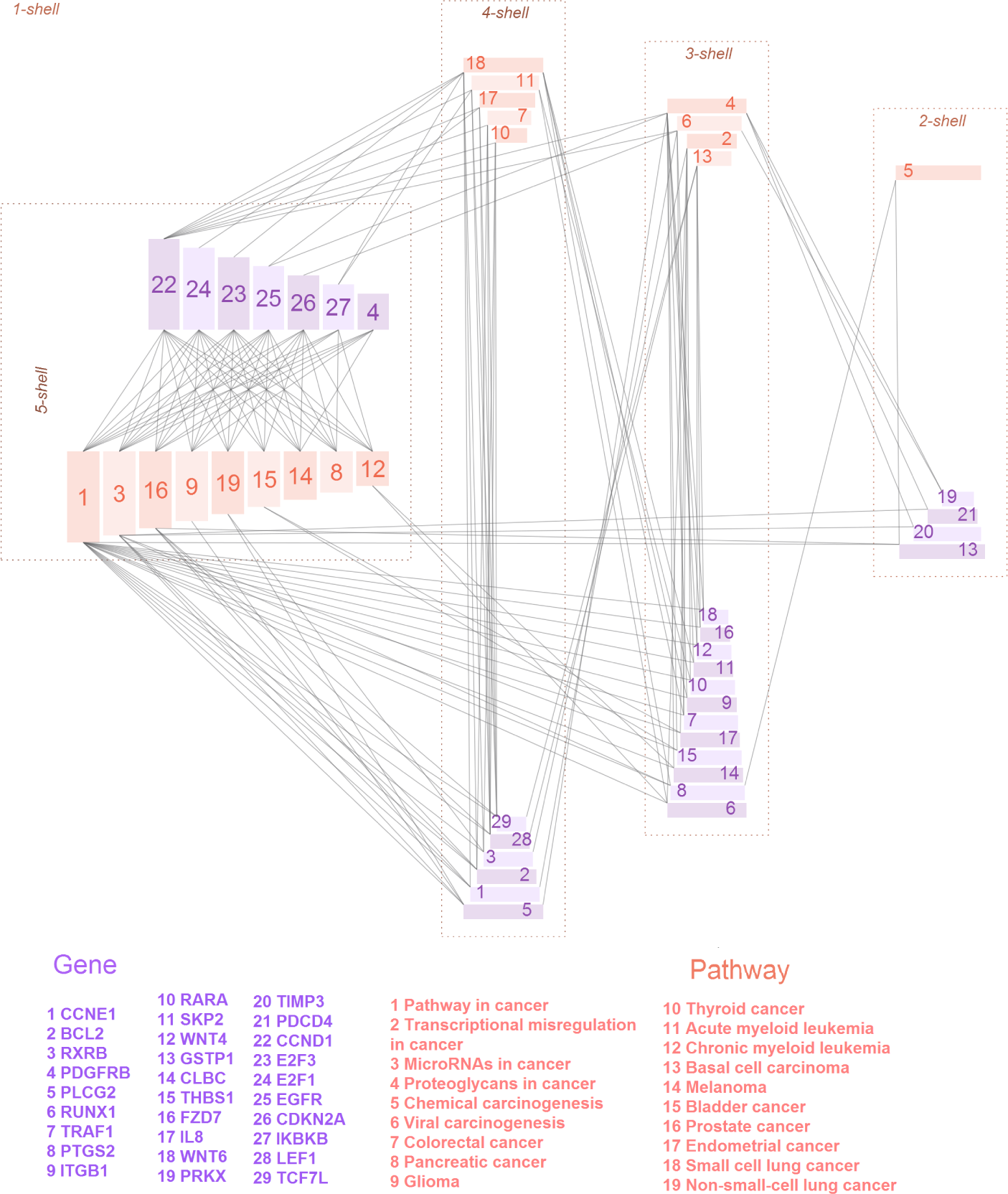
Gene-pathway association network [41].

Figure 6 shows a network with a stronger hierarchy than figure 5. The identification of genes most frequently associated with pathways to cancer is straightforward.

The main application of the polar plot is the visual comparison of networks even if their sizes are very different. Figure 7 is a subset of a disease-cofactor network. Authors selected diseases tied to at least 5 cofactor-interacting proteins (39 nodes) and plotted the bipartite graph 7. The ziggurat plot (fig. 7A) of the subset shows an extremely nested structure, an effect of the selection rule. The polar plot of this network fragment (fig. 7B) has an uncommon organization of disease nodes, at almost the same *k_radius_* distance of the center. The polar plot of the full network (fig. 7B), with 414 nodes, displays a much richer structure. Diseases are distributed across a wider range of *k_radius_*. Most cofactors have high degree and were not filtered. As a result, the structure of this guild is very similar in both polar plots.

**Figure 7:**
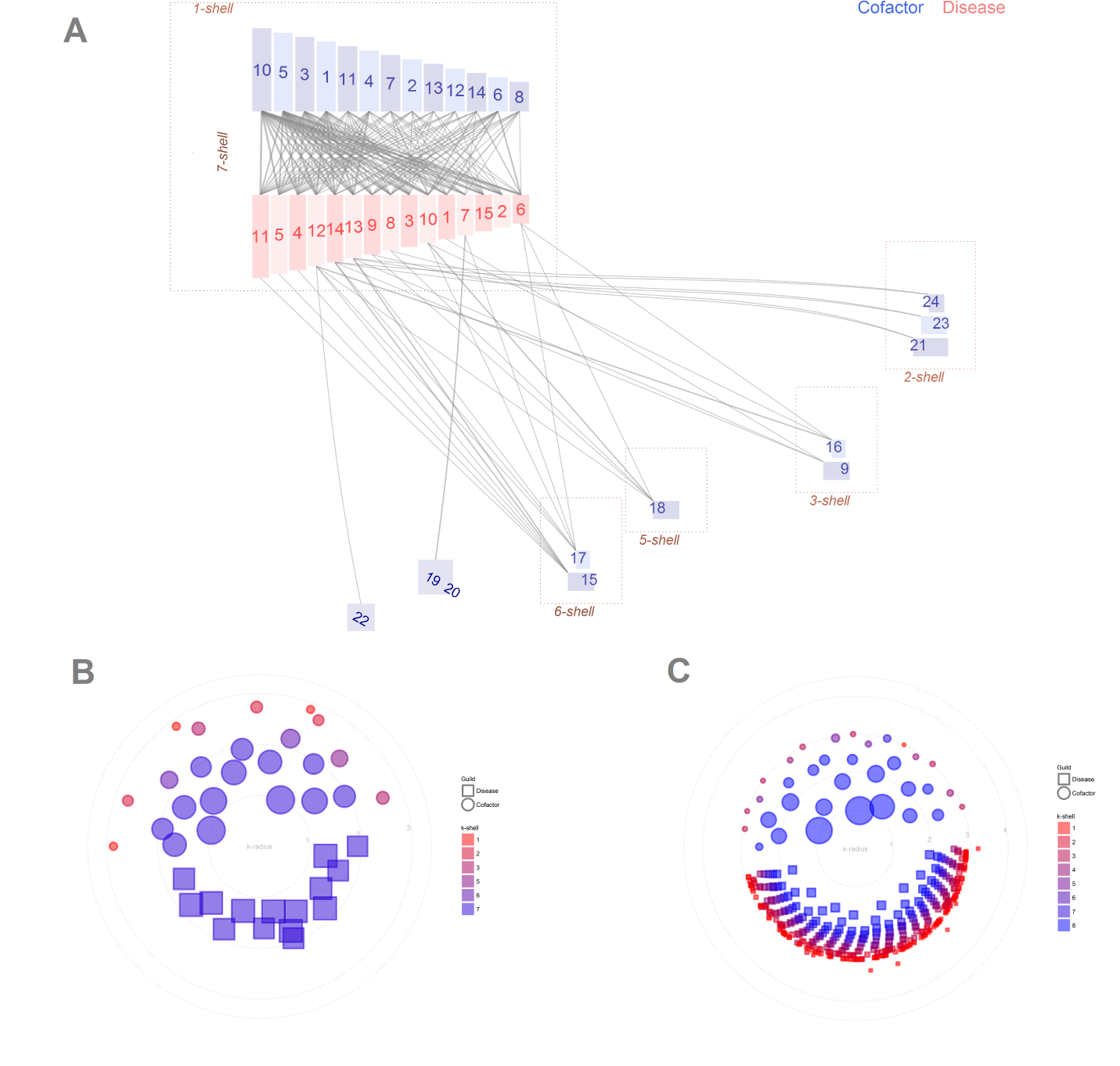
Disease-cofactor network [42]. A: ziggurat plot of the subset. B: polar plot of the subset. C: polar plot of the full network.

These figures are a small sample of the importance of choosing a good visualization tool with a correct analysis of decomposition of a network.

## 4. Software

The *k-core* analysis and plotting of ziggurat and polar graphs is provided as an open source application.

### 4.1. The kcorebip package

The `R` package `kcorebip` contains the functions to perform the analysis and to plot static graphs of a network. It comes with a set of networks for testing purposes. Ecological data were downloaded from the *web of life* database [43]. As the format of the *web of life* files has became a standard *de facto* by its simplicity, `kcorebip` follows the same convention for input files.

The function `network k analysis` computes the *k-magnitudes* and other useful indexes, using the functions that provide packages as `bipartite` and `igraph` [44, 45]. We refer to the user manual for details.

Ziggurat and polar graphs use basic calls to the `ggplot2` graphics package [46]. We compute from scratch coordinates and sizes, not relying on other network plotting libraries.

### 4.2. Interactive application

The `kcorebip` package is a powerful solution for researchers with programming skills that need high quality plots for scientific publications, but exploratory analysis requires a more interactive approach. `Bipartgraph` has designed with this need in mind.

The technological choice is `Shiny`, the `R` reactive programming environment. It has the advantage of a native backend and a `Javascript`-based user interface that may be easily extended. This combination of technologies ensures a wide compatibility with most common operating systems.

The `Interactive Ziggurat` is the main feature of `Bipartgraph`. The original implementation of the `kcorebip` package only provided the `ggplot2` object to display or save. To create an interactive version we faced two main choices, replicating the code with a dynamic technology or extending `kcorebip`. We found a fast and almost non intrusive solution creating an `SVG` object. The ziggurat is a set of rectangles, lines and texts. The most time consuming tasks are network analysis and spatial distribution. These computations are performed just once, and besides each `ggplot2` element the function plots, it creates the `SVG` equivalent.

The browser displays the `SVG` ziggurat with multiple options for the user: tooltips, select a node or a link, highlight connections, zoom in and zoom out. In addition, a second panel shows information of highlighted nodes and the available information on Wikipedia (fig. 8).

**Figure 8:**
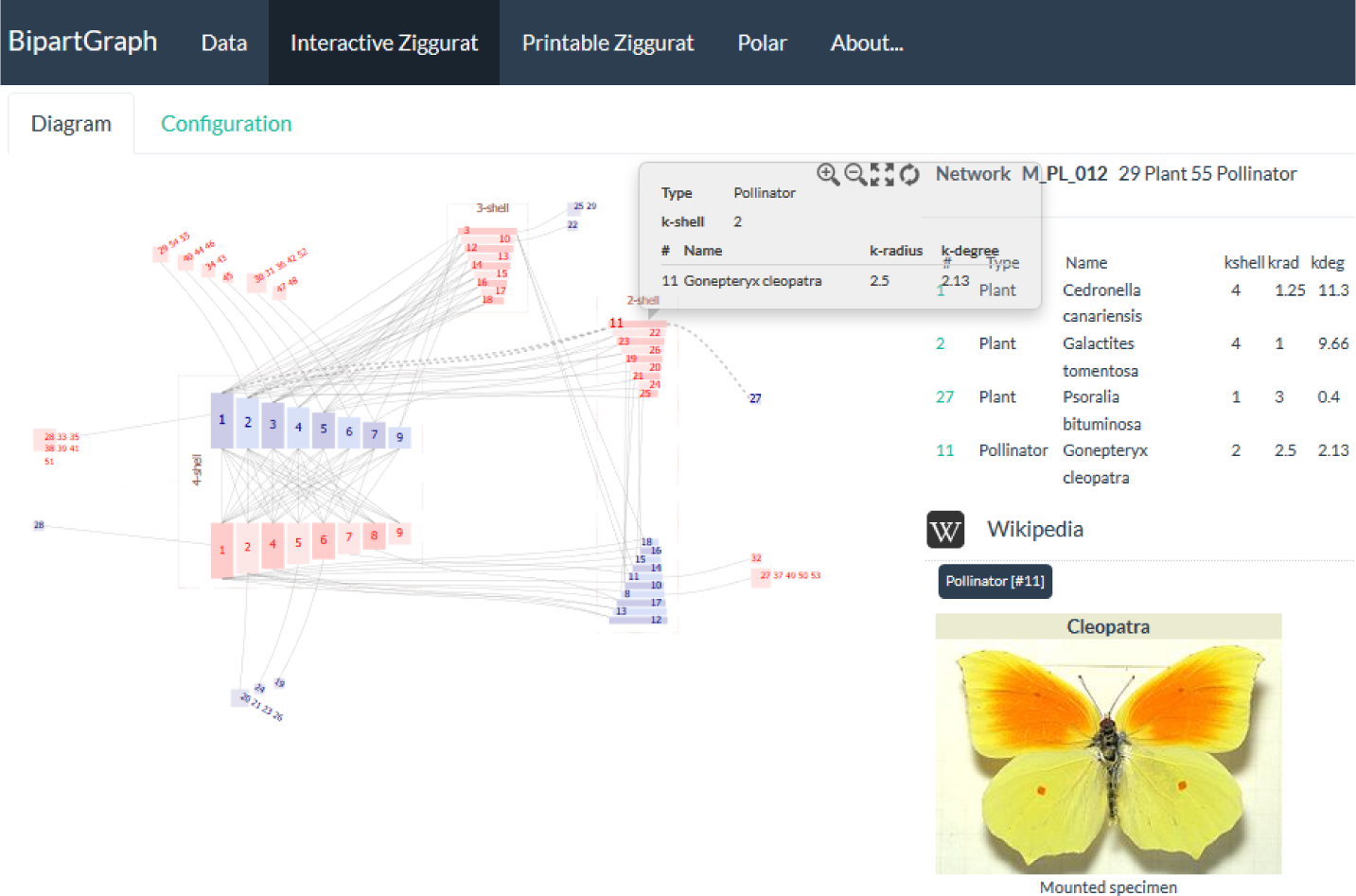
Interactive ziggurat user interface. Plant-pollinator assembly in Garajonay, Canary Islands (Spain), recorded by Olesen. Highlighted the pollinator species number 11,*Gonexpteryx cleopatra*

The configuration panels make plot properties easy to modify. Visual and intuitive Shiny controls, as sliders or checkboxes, hide the complexity of the input parameters of the `ziggurat graph` function.

At any moment, the user may download the high quality, high resolution static plot with the `Printable Ziggurat` option. In order to reproduce the results or to include the graph in other environments, such as `R Markdown` or `Jupyter` notebooks, we added the `Download generating code` button. When clicked upon, `Bipartgraph` writes a file with the last `ziggurat graph` call, ready to use in any `R` script.

There is not interactive version of the `Polar` plot, as we think that network exploration is much easier with the ziggurat. The user may produce the static polar plot, the high quality downloadable `PNG` file and the generating code, in the same way that we have explained with the ziggurat.

## 5. Conclusions

Visualization of bipartite biological networks is very useful for researchers when they are interested in following the paths from a node or scanning the structure of the network. Using the *k-core* decomposition we have designed and developed two new graphs that work by information reduction (Polar plot) and spatial grouping by connectivity (Ziggurat plot). They provide two complementary views of internal network structure.

We would like to emphasize the importance of choosing a correct visualization of complex networks, and in particular of bipartite networks, that helps a correct understanding of networks of a large number of nodes and high density.

Software is provided as Open Source, under a very loose MIT license, and comes in two versions. The package kcorebip provides the full functionality for researchers with a minimum of R programming skills. The application Bipartgraph is the fully fledged interactive environment to build both kind of graphs for this public. Its user centric design makes it very easy to master, provides some additional features and is open to new fields of application such as education.

## Software Availability

Name of software: BipartGraph

Programming language: R

Operating system: Windows, Linux and MacOS

Availability: SW at https://github.com/jgalgarra/bipartgraph

User interface: Web browser

License: Free, under MIT License

## Acknowledgements

This work was supported by Ministry of Economy and Competitiveness of Spain (MTM2015-63914-9).

## Data availability

Interaction matrixes were downloaded from the web of life database http://www.web-of-life.es/. A subset of these matrixes is installed by default with*BipartGraph*, including all networks used in this paper.

